# Maternal Immunoglobulin A regulates the development of the neonatal microbiota and intestinal microbiota-specific CD4+ T cell responses

**DOI:** 10.1101/2024.06.10.598156

**Authors:** Darryl A. Abbott, Ali T. Rai, Aaron Yang, Yixuan Cai, Shelcie Fabre, Austin J. Frazer, Jacob D. Deschepper, Amanda C. Poholek, Timothy W. Hand

## Abstract

Breast milk is a complex mixture of nutrients and bioactives that promote infant development and decrease the incidence of chronic inflammatory disease. We investigated the role of one milk-derived bioactive, Immunoglobulin A (IgA) on the developing small intestinal microbiota and immune system. We demonstrate that early in life, milk-derived IgA suppressed colonization of the small intestine by *Enterobacteriaceae* and regulated the maturation of the small intestinal epithelium and the development of intestinal IL-17-producing CD4^+^ T cells. *Enterobacteriaceae*- specific CD4^+^ T cells, induced in the first weeks of life in the absence of milk-derived IgA, persisted in the intestine as memory T cells that can contribute to inflammatory disease later in life. Our study suggests that milk-derived IgA shapes mucosal immunity by regulating the neonatal microbiota thus preventing the development of long-lived intestinal microbiota-specific T cells.

## Introduction

The assembly of the intestinal microbiome begins at birth and the basis for the adult bacterial community is formed in the first years of life (*1, 2*). Because the microbiome is acquired from the environment, the timing and magnitude of intestinal colonization by different bacterial taxa is important to host health, in both the near and long-term (*3*). Disruption of the microbiome during the neonatal period, through infection or antibiotic use, is a risk factor for autoimmunity and immunopathology(*4–6*), and suggests a window of opportunity for establishment of host immunity via a regulated acquisition of the microbiota (*7*).

Breastmilk contains a variety of nutrients and bioactive molecules that support infant health in both the short- and long-term and breastfeeding is associated with reduced childhood infection and a decreased incidence of allergy, obesity, and autoimmunity(*8–10*). As infants transition from sterility to ubiquitous microbial colonization, they are particularly vulnerable to infection, because they lack the protection provided by immune memory (*11, 12*). To counteract this vulnerability, breastmilk contains bioactive molecules that regulate intestinal bacteria, such as antibodies that transfer immune memory from the mother to the infant (*8, 13*). In addition to protection from pathogens, breastmilk-derived antibodies are important in shaping the development of the neonatal microbiota and immune system(*14–18*).

Dimeric secretory IgA (sIgA), is produced in prodigious amounts at mucosal sites where it protects the host through diverse non-cytolytic effects on pathogenic and commensal bacteria (*19–21*). SIgA is transported across the epithelium by polymeric immunoglobulin receptor (PigR), which remains bound to IgA as secretory factor and increases its half-life in the intestine by reducing proteolytic cleavage (*22*). SIgA is the predominant antibody (>90%) secreted into breastmilk and during the first month of life in rodents and humans, milk is the only source of IgA(*11, 12*). IgA-producing plasma cells traffic from the Peyer’s Patches and intestine to the mammary glands during pregnancy (*23–26*). The anti-bacterial reactivity of breast milk IgA is highly heterogeneous between individuals but stable over time, which highlights a drawback of ‘vertically transmitted’ immunity, that it is not responsive to the conditions of the infant’s intestine(*27*). There are potential consequences for the infant of ‘holes’ in the anti-bacterial antibody repertoire, most notably an increased susceptibility to enteric pathogens(*17, 18, 28*) and a reduction in IgA binding to *Enterobacteriaceae* is associated with the development of Necrotizing Enterocolitis in preterm infants(*11*).

To investigate how milk-derived maternal IgA (matIgA) shapes the development of the neonatal microbiome, intestine and immune system we developed a breeding system where genotypically identical immunocompetent IgA^+/-^ mice pups were nursed by either IgA-deficient or sufficient dams. Using this system, we discovered that matIgA was critical to the development of the neonatal microbiota and microbiota-responsive intestinal CD4^+^ Helper T cells. Critically, CD4^+^ T cells activated during the neonatal period survived into adulthood as memory T cells where they can affect host/microbiota interaction long-term.

## Results

### Maternal IgA regulates the assembly of the neonatal small intestinal microbiota

To isolate the effects of matIgA on the development of the neonatal microbiota and intestinal immune system we developed a heterozygous breeding scheme (**Figure 1A**) which enables the comparison of genetically identical and immunocompetent (IgA^+/^) mouse pups nursed by either an IgA^-/-^ dam or an IgA-sufficient C57BL/6 dam. To create controls, C57BL/6 dams are bred to IgA^-/-^ sires, which ensures that the atypical microbiota of IgA^-/-^ mice(*29*) is present in each cage. We confirmed that pups nursed by IgA^-/-^ dams have minimal IgA binding on their intestinal bacteria prior to weaning (**Figure 1B-C**), indicating that almost all intestinal IgA in the pre-weaning period is derived from the dam’s milk (*11, 14*). As previously described, in the absence of maternal IgA (matIgA) there is a compensatory increase in IgM and IgG bound bacteria(*30, 31*) (**Figure 1B**), but this modest increase remains about ½ the total amount of bacteria bound by IgA in pups of C57BL/6 dams. Thus, maternal IgA deficiency both eliminates pre-weaning neonatal intestinal IgA and reduces the total amount of antibody bound to intestinal bacteria.

**Figure 1.**
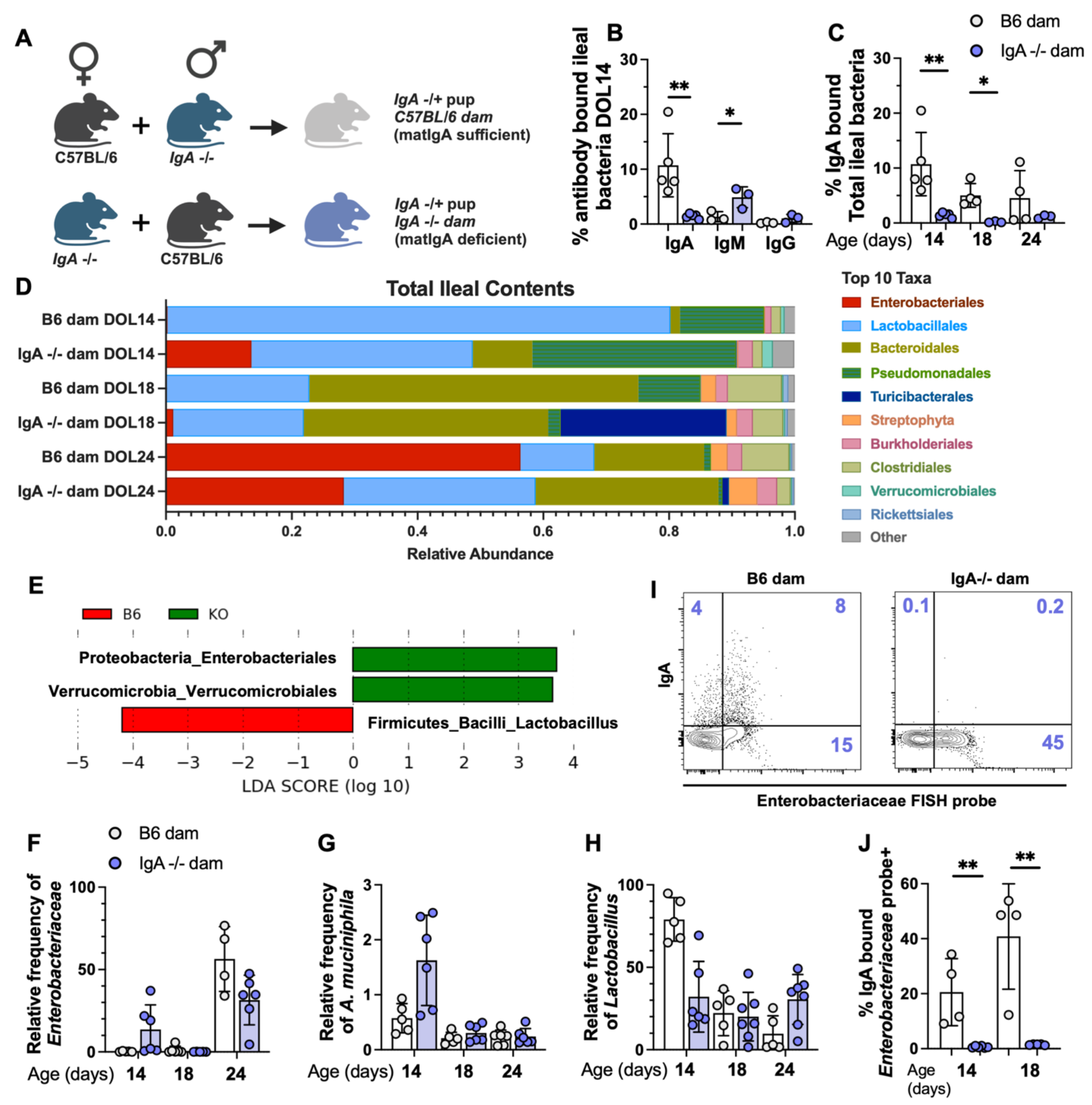
– Maternal IgA regulates the colonization of the infant small intestine by the microbiota. (A) Mice were bred such that all pups would be IgA+/- and differ in the IgA production status of the dam (nursed by B6/IgA+/+, grey circles; nursed by IgA-/-, blue circles). (B) Percent antibody (IgA, IgM or IgG) bound Hoescht+ (DNA+) events within total ileal contents (TIC) of DOL14 pups. (C) Percent IgA bound Hoescht+ (DNA+) events in TIC of IgA +/- pups at DOL 14, 18, 24. (D-H) Genomic DNA isolated from TIC of pups bred as described in (A). TIC analyzed by 16S rRNA gene amplicon sequencing. Samples were collected from ≥2 separate cages for each group to minimize cage effects, with a minimum of 4 animals per experimental group. (D) Mean relative abundance of top 10 bacterial orders. (E) LEfSe analysis of (D) at DOL14; taxa shown are significantly different (LDA score (log10)>2) between C57BL/6 (B6) or IgA -/- (KO) fed pups. (F-H) relative abundance of selected bacterial taxa. (I) Flow cytometric analysis of TIC (DOL14) by fluorescent in situ hybridization (FISH) of pups fed by either IgA+/+ or IgA -/- dams. (J) Quantification of (I). *Enterobacteriaceae* IgA± events in (I-J) are pregated on Hoescht+ (DNA+) *Eubacteria*+ (Eub338). Data represent 1 (D-H) and 3 (B-C, I-J) independent experiments with at least 3 samples per group. Unpaired student’s T test was used for all analysis. *p<0.05, **p<0.01, ***p<0.001. Each point represents one mouse. (A) created on BioRender.

We next measured the effect of matIgA on the small intestinal microbiota by 16S rRNA gene amplicon sequencing of samples that spanned the nursing and weaning process. Principal coordinate analysis of β diversity revealed time post birth as the primary differential separating samples (**Figure S1A-C**). There were also differences associated to the IgA status of the dam. Distinctly, in day-of-life (DOL) 14 ileal samples (**Figure 1D-E**) there was a significant increase in the fraction of *Enterobacteriaceae* and *Akkermansia* in the small intestine of pups nursed by IgA^-/-^ dams and a corresponding decrease in *Lactobacillus* (**Figure 1E-H**). *Lactobacilli* are the putative metabolizers of milk oligosaccharides in mice, so their replacement could be important to the pups(*3*). The differences in the structure of the microbiota between pups of IgA^-/-^ and control C57BL/6 dams were much less apparent at DOL18, likely due to the dominant effect of switching to solid food over any possible effects of matIgA (**Figure 1D** and **S1A**). Similar to DOL14, at DOL24, *Enterobacteriales* was amongst the most common taxa, but did not differ according to milk IgA status. One explanation for the difference between DOL14 and DOL24 is that by the later timepoint pups have largely self-weaned but are themselves making very little IgA (*14*), equalizing the intestinal IgA levels amongst the pups.

In both mouse models and adult humans, intestinal *Enterobacteriaceae* bacteria are enriched for IgA binding, and IgA deficiency increases their relative abundance (*30–32*). Using Fluorescence In Situ Hybridization (FISH) in combination with flow cytometry we discovered that approximately 20-40% of *Enterobacteriaceae* cells were bound by IgA in the preweaning intestine of C57BL/6 pups (**Figure 1I-J**), which is a substantial enrichment over the IgA binding of the total microbiota (∼5-10% IgA bound). Taken together, we have demonstrated that matIgA preferentially binds *Enterobacteriaceae* and that these bacteria are significantly expanded in pups if matIgA is absent.

### Development of the neonatal small intestinal epithelium is regulated by maternal IgA

We next analyzed the primary contact point between the host and microbiota, the small intestinal epithelium. Principal coordinate analysis of flow cytometrically isolated (Epcam^+^CD45^-^; **Figure S2A**) epithelial cell transcriptomes (RNAseq) revealed that DOL14 samples clustered away from DOL24 and DOL42, indicating that this timepoint of development represents a distinct developmental window (**Figure S2B**). In concert with our microbiota data (**Figure 1**) clear differences between pups nursed by IgA^-/-^ and C57BL/6 dams were only observed at DOL14.

The small intestine is composed of distinct epithelial cell subtypes within two lineages; absorptive enterocytes and secretory cells (Enteroendocrine, Goblet, Paneth and Tuft) that differentiate from the recent daughter cells of Intestinal Stem Cells (ISCs). Analysis of the genes most significantly different between IgA^-/-^ and C57BL/6 fed pups at DOL14 (**Table S1**) revealed canonical genes (Atoh1, Muc2, Chga, Dckl1, Reg3b) associated with the various secretory cell types were enriched in IgA^-/-^ fed pups. Hierarchical clustering and GSEA analysis revealed that the gene programs associated with secretory cells, but not the absorptive or ISC cell types, were enriched at DOL14 in the absence of matIgA (**Figure S2D-E**). Microscopic quantification of small intestinal enteroendocrine (CHGA^+^) and goblet (Wheat Germ Agglutinin^+^) cells showed significant increases in the absence of matIgA at DOL 14, but no changes observed at DOL28 (**Figure S2F-H**). *Akkermansia muciniphila*, which we observe to be increased in the absence of matIgA, can directly induce the differentiation of intestinal epithelial cells to Paneth and goblet cells by acting directly on LGR5+ stem cells(*33*). Thus, during the critical first weeks of life when IgA is not being made by neonates, matIgA regulates both the development of the microbiota and the small intestinal epithelium.

### Maternal IgA regulates the development of small intestinal T Helper 17 T cells

The acquisition of the microbiome is a critical regulator of multiple aspects of the intestinal immune response(*6, 34–36*). MatIgA regulation of the microbiota impacts the development of colonic regulatory T cells (Tregs) and matIgA-dependent changes to immunity can be transmitted into adulthood or even into the next generation(*7, 35, 37*). The role of matIgA in modulating effector T cell activation and differentiation in the neonatal small intestine is less clear. We quantified activated T helper cells and Tregs in the small intestinal lamina propria (siLP), intestinal epithelium (IEL), Peyer’s Patches (PP) and mesenteric lymph node (mLN) at DOL 24, 6-10 days following the period when we observed shifts in the microbiota and epithelial cells, and when putative matIgA-dependent T cell responses should be at their peak. The number and frequency of colonic RORγt^+^ Tregs is regulated by matIgA binding of bacteria in the neonatal intestine (*37*). Although the total number of siLP Tregs were not significantly altered by a lack of matIgA (**Figure 2A** and **S3A**), we did observe an increase in the number siLP RORγt^+^ FoxP3^+^ T cells (**Figure 2B** and **S3B**). Measurement of the number of CD4^+^ effector T cells (CD4^+^CD44^hi^FOXP3^-^) in the small intestine and associated lymphoid tissue also revealed a substantial (∼10x) increase amongst siLP cells (**Figure 2C**). Further characterization of siLP CD4^+^ Effector T cells revealed significant increases in both RORγt^+^ IL-17 expressing Th17 cells and T-bet^+^ IFNγ-expressing Th1 cells with similar but modest differences observed in intestine associated tissue sites (**Figure 2D-F** and **S3C-E**). In the pups of matIgA deficient dams, we also observed an increase in Th17 cells that co-express IFNγ, which have been shown to be more pathogenic in a variety of disease settings (**Figure 2F**) (*38, 39*). We observed a similar increase in the proportion of Th17 (CD4^+^RORγt^+^) in the siLP of pups when using either Rag1^-/-^ knockout dams, which make no T and B cells or ‘IgMi’ dams(*40*), which are unable to secrete antibodies, indicating our findings are not unique to the IgA^-/-^ model (**Figure S3F**).

**Figure 2.**
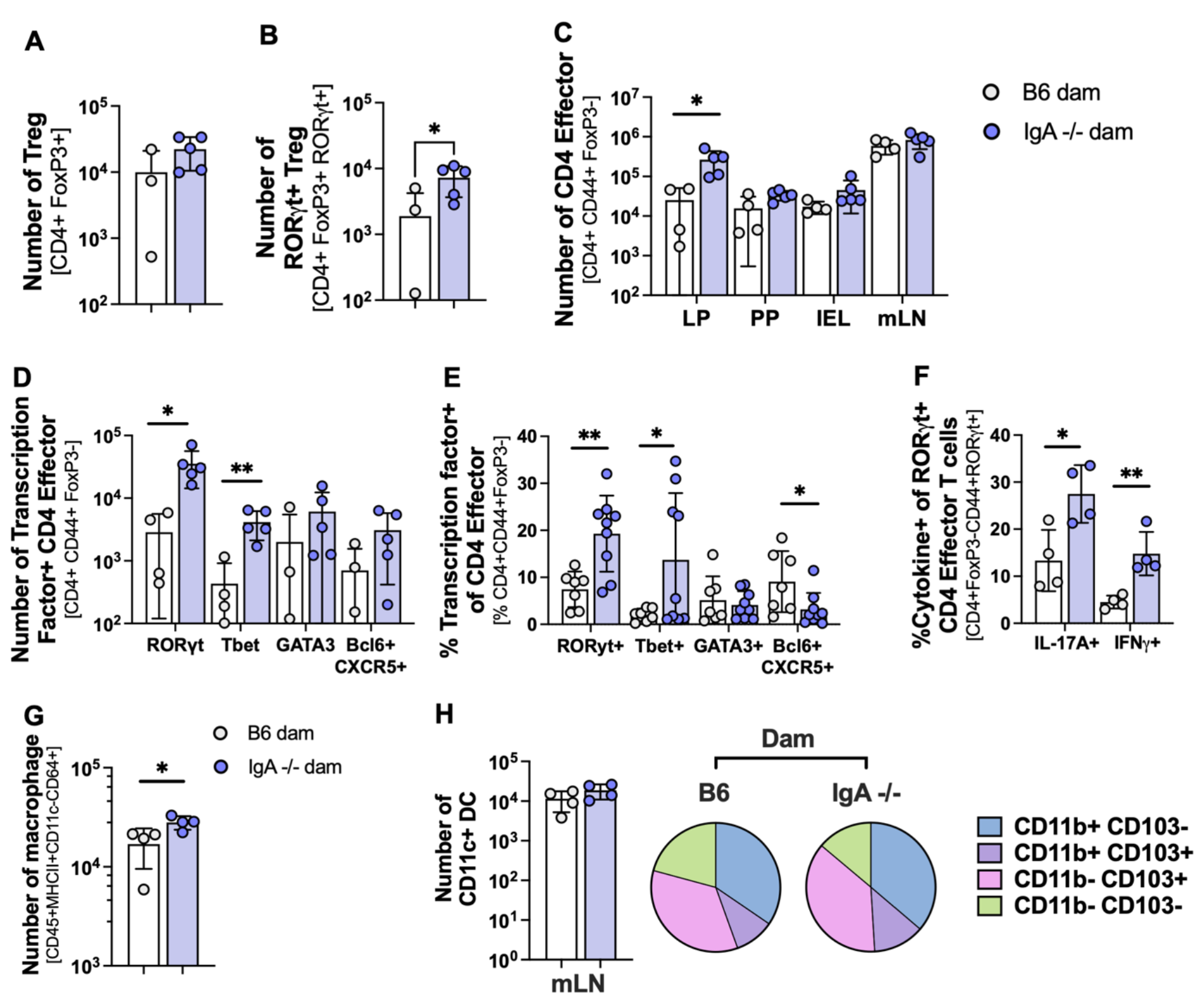
- Maternal IgA regulates Th17 cell activation and accumulation in the neonatal small intestine. Pups were bred as in Figure 1A (nursed by B6/IgA+/+, grey circles; nursed by IgA-/-, blue circles). (A-G) Flow cytometric analysis of pups from DOL24. (A) Total count of Tregs (Live TCRβ+CD4+CD90+Foxp3+) in the siLP. (B) Total count of peripheral (Rorγt+) Tregs (Live TCRβ+CD4+CD90+Foxp3+ RORγt) in the siLP. (C) Total count of T effector cells (Live TCRβ+CD4+CD90+Foxp3-) in indicated tissues. (D) Total count and (E) Relative proportion of T effector cells in the siLP positive for indicated transcription factors (Tfh = CXCR5+ Bcl6+). (F) Proportion of RORγt+ cells in the lamina propria expressing the indicated cytokines following PMA/Ionomycin stimulation. (G+H) Flow cytometric analysis of pups from DOL14. (G) Total count of macrophages (Live CD45.2+MHCII+CD11c-CD11b+CD64+) in the siLP. (H) Total count of mLN dendritic cells (DC) (Live CD45.2+MHCII+CD64-CD11c+). Pie chart of averaged CD103 and CD11b expression within DC subsets in IgA +/- pups fed by either a B6 or IgA -/- dam as indicated. Unpaired student’s T test was used for all analysis. *p<0.05, **p<0.01, ***p<0.001. Data representative of 2-4 independent experiments with 3-6 mice per group. Each point represents one mouse.

After birth intestinal myeloid cells develop along a characteristic trajectory(*3, 41*). We hypothesized that differences in antigen-presenting dendritic cells or macrophages might be directing differences in T cell differentiation. In the absence of matIgA, there was a modest but significant increase in the number of siLP CD64^+^ macrophages (**Figure 2G**), which can support Th17 T cell differentiation in the LP(*42, 43*). Measurements of siLP and mLN dendritic cell subsets in pups of IgA deficient and sufficient dams, revealed no significant differences, including in the CD103^+^ CD11b^+^ DCs that have been associated with Th17 induction(*44*) (**Figure 2H** and **S3G)**. Innate Lymphoid cells are also important regulators of neonatal Tregs(*45–47*) and can also be affected by the early life microbiota/maternal effects(*48*), but we observed only modest differences in siLP ILC2 frequenices and no differences in ILC numbers associated with the presence or absence of matIgA (**Figure S3H-I**).

### Induction of neonatal Th17 T cells in the pups of IgA^-/-^ dams is dependent upon the microbiota

Intestinal Th17 cells are induced in response to microbiota-derived antigens and metabolites(*49*). Therefore, we tested whether matIgA regulation of the neonatal microbiota (**Figure 1**) is responsible for limiting the activation of Th17 T cells in the intestine. We first assayed whether the neonatal microbiome was sufficient to induce Th17 responses in the small intestine by colonizing adult germ-free mice with the DOL14 small intestinal contents of pups nursed by either a C57BL/6 or an IgA -/- dam (**Figure 3A**). Consistent with findings in pups of IgA-/- dams, colonization of germ-free pups with the small intestinal microbiota of pups nursed by IgA deficient dams and not controls had substantially increased populations of intestinal but not mLN CD4^+^ RORγt^+^ Th17^+^ T cells (**Figure 3B-D**).

**Figure 3.**
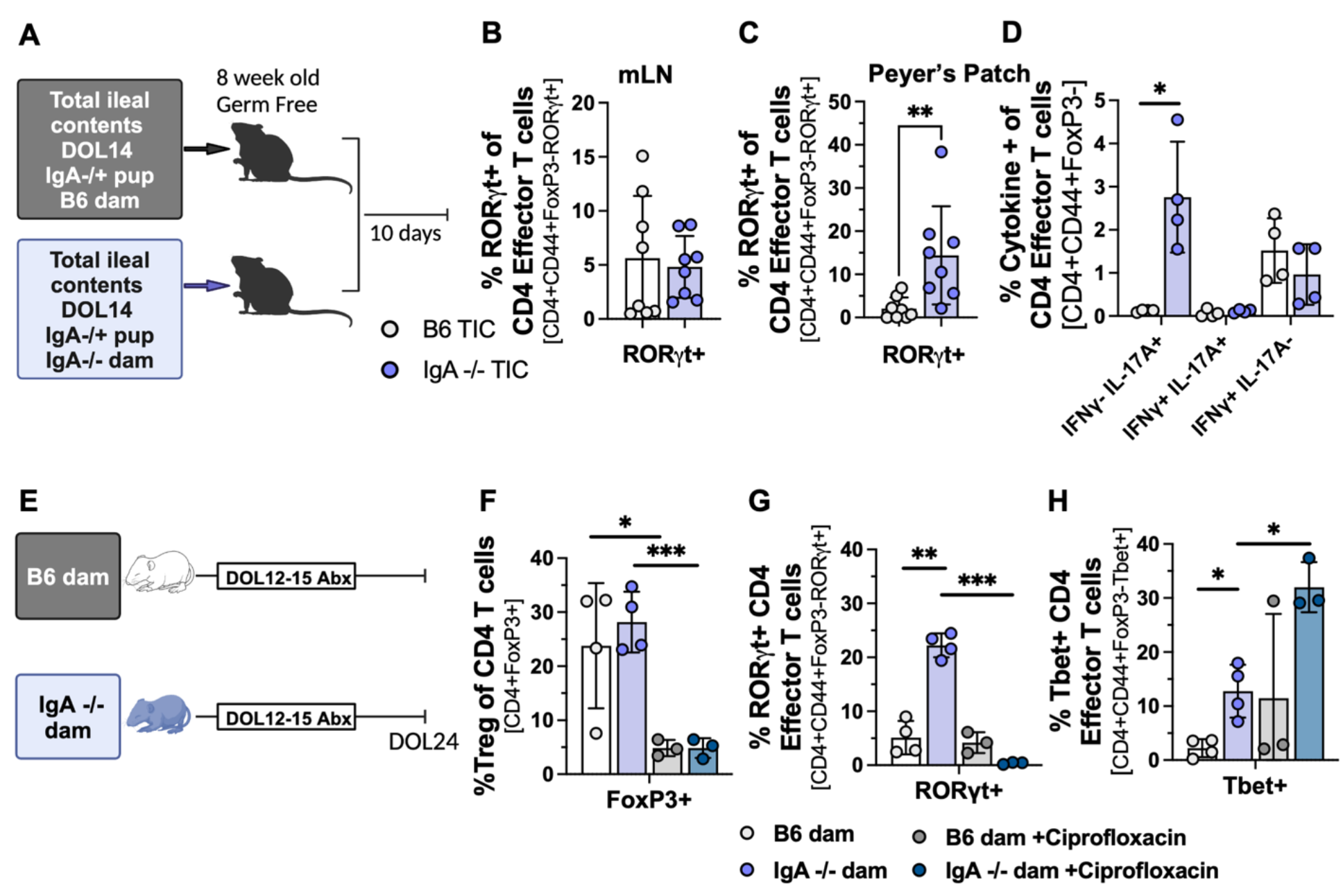
- Induction of neonatal Th17 T cells in the pups of IgA^-/-^ dams is dependent upon the microbiota. Pups were bred as in Figure 1A. (A) Fecal microbiota transplant (FMT) of the total ileal contents of DOL14 pups (grey circles = B6/IgA+/+ fed; blue circles = IgA-/- fed) into Germ-free mice. (B-D) 10 days post FMT, T cells were isolated from mLN and PP and assessed by flow cytometry. (B) Relative proportion of T effector (Live TCRβ+CD4+CD90+Foxp3) cells expressing RORγt in mLN (C) Relative proportion of T effector (Live TCRβ+CD4+CD90+ Foxp3+) cells expressing RORγt in the PP (D) Relative proportion of T effector cells from the PP producing indicated cytokines following PMA/Ionomycin stimulation. (E) Pups bred as in Figure 1A were treated with Ciprofloxacin by gavage (DOL12-14). (F-H) Flow cytometric analysis of the relative proportion of T effector cells from the siLP at DOL24 expressing FoxP3 (F), RORγt (G), and Tbet (H). Unpaired student’s T test was used for all analysis. *p<0.05, **p<0.01, ***p<0.001. Data representative of two independent experiments with 3-6 mice per group. Each point represents one mouse. A and E created on Biorender.

Analysis of the small intestinal microbiota at DOL14, indicated that in the absence of matIgA pups had significantly increased *Enterobacteriaceae* (**Fig 1E**), intestinal *Enterobacteriaceae* is enriched for binding by matIgA (**Figure 1J**) and some *Enterobacteriaceae* bacteria are sufficient to induce intestinal Th17 responses(*50*). To target *Enterobacteriaceae* in the intestine we used ciprofloxacin, a fluorinated quinolone antibiotic which is effective against *Enterobacteriaceae* but far less effective against the other bacteria (*Akkermansia*, *Lactobacillus*) that dominate the neonatal mouse intestine(*51, 52*). To test whether *Enterobacteriaceae* is necessary for the neonatal induction of CD4^+^ Th17 T cells in the absence of matIgA, we gavaged pups from DOL12 to DOL15 with ciprofloxacin which eliminated intestinal *Enterobacteriaceae* (data not shown) and reduced small intestinal FOXP3+ Tregs (**Figure 3E-F**). Critically, ciprofloxacin treatment diminished the difference in siLP Th17 cells between IgA-/- and C57BL/6-fed pups at DOL24 (**Figure 3G**). In contrast, Tbet expressing T cells were significantly increased (**Figure 3H**) indicating that matIgA dependent increases in these cells are not regulated by the microbiota. Taken together, our results indicate that matIgA regulates the assembly of the neonatal microbiota, in particular *Enterobacteriaceae*, and that failure of matIgA microbial regulation leads to the early activation of IL-17-producing intestinal T cells.

### Neonatal microbiota specific T cells activated in the absence of maternal IgA persist into adulthood

Infants fed formula, which lacks IgA, have significantly higher incidence of many autoinflammatory diseases, including Inflammatory Bowel Disease(56). One hypothesis consistent with this finding would be that T cells activated in the absence of matIgA populate the host tissues and in instances of reduced barrier function, are re-activated and contribute to inflammation. To test this possibility, at 8 weeks post-birth we gavaged mice with indomethacin, a non-steroidal anti-inflammatory drug (NSAID), that induces small intestinal damage in an IL- 17A-dependent manner and increases intestinal *Enterobacteriaceae* (*53, 54*). Indomethacin treatment of eight-week-old mice nursed by IgA^-/-^ dams revealed increased inflammation compared to controls, as indicated by increased weight loss (**Figure 4A** and **S4A**). Furthermore, antibody-mediated depletion of CD4^+^ T cells during indomethacin treatment demonstrated that CD4 T cells are necessary to drive the weight loss in the adult offspring of IgA^-/-^ dams (**Figure 4A**). Thus, five weeks post weaning, genotypically identical mice that differ only in the IgA status of their dam, produce different responses to small intestinal damage that are dependent upon the presence of CD4^+^ T cells.

**Figure 4.**
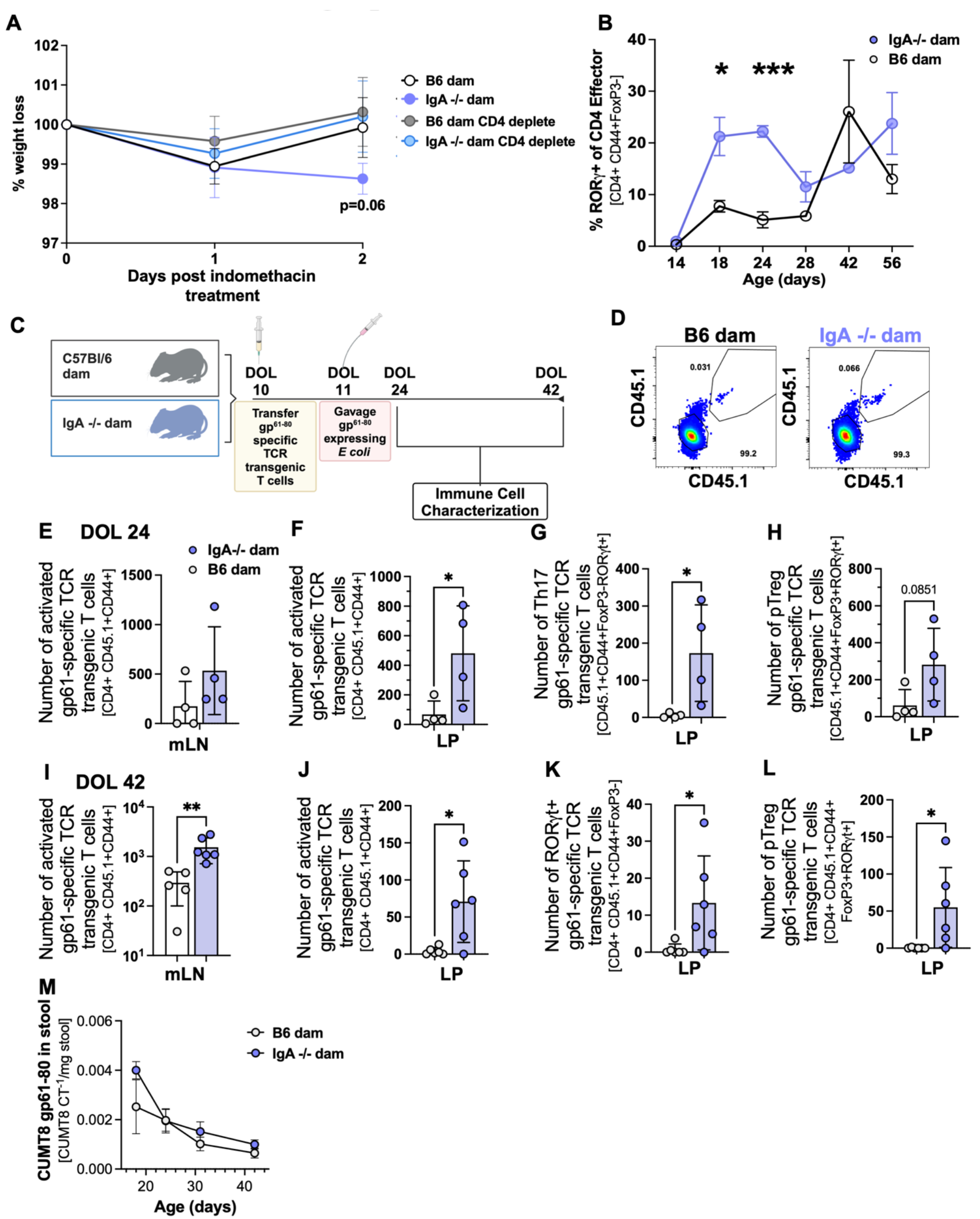
Neonatal microbiota specific T cells activated in the absence of maternal IgA persist into adulthood. Pups were bred as in Figure 1A. (A) IgA +/- pups were cohoused following weaning until eight weeks of age, at which point all groups were gavaged with 6.25mg/kg/day of indomethacin for three days. Half of mice were treated on day -3 and 0 with anti-CD4 antibodies. Weight was monitored daily. Data is pooled from 2 independent experiments with 3-6 mice per group. (B) Longitudinal analysis total RORγt+ cells in the siLP of pups bred as in Figure 1A . Graph depicts cells gated (Live TCRβ+ CD4+ FoxP3-RORγt+). Data representative of 2-4 independent experiments for each time point. (C) 1x10^5^ CD45.1+ naïve (CD44^lo^CD25-) GP_61-80_ TCR transgenic T cells were transferred by IP injection into pups at DOL10-11. CUMT8 GP_61-80_ was gavaged one day after transfer. Lamina propria and mLN from mice in (C) were assessed post transfer at DOL24 (D-H) and DOL42 (I-L). (D) Flow cytometric plot of mLN CD45.1+ GP_61-80_ TCR transgenic T cells cells at DOL24. Gated on Live TCRβ+CD4+. (E-H) Analysis of CD45.1+ CD4+ T cells isolated from DOL24 IgA +/- pups. (E) Total count of activated CD45.1+ CD4+CD44+ T cells in the mLN. (F) Total count of activated CD45.1+ CD4+CD44+ T cells cells in the siLP. (G) Total count of CD45.1+ CD4+ T cells expressing RORγt+ in the siLP. (H) Total count of CD45.1+ CD4+ T cells expressing RORγt+FoxP3+ in the siLP. (I-L) Analysis of CD45.1+ CD4+ T cells cells isolated from DOL42 IgA +/- pups. (I) Total count of activated CD45.1+ CD4+ CD44+ T cells cells in the mLN. (J) Total count of activated CD45.1+ CD4+CD44+ T cells in the siLP. (K) Total count of CD45.1+ CD4+ T cells expressing RORγt+ in the siLP. (L) Total count of CD45.1+ CD4+ T cells expressing RORγt+FoxP3+ in the siLP. (M) DNA extracted from stool following CUMT8 gavage on day 10 was assessed weekly until day 42 for CUMT8 flic3 transcript by qPCR. The inverse of three technical replicates average CT value was divided by stool weight. (K) Unpaired student’s T test was used for analysis of (A- I) and simple linear regression for (E). *p<0.05, **p<0.01, ***p<0.001. Data representative of two independent experiments with 3-6 mice per group. Each point represents one mouse. C was created with BioRender

We next wanted to determine the longevity of CD4^+^ T cells activated during the neonatal period in the absence of matIgA. Longitudinal measurements of pups nursed by IgA deficient dams and controls reveals that the time period when we can measure increased RORγt^+^ Th17 cells in the siLP is limited to the first four weeks of life (**Figure 4B**). However, this result does not exclude the possibility that microbiota-specific CD4^+^ T cells can survive as memory cells. To track microbiota-specific CD4^+^ T cells we modified an adherent-invasive strain of *Escherichia coli* (CUMT8)(*54*) to express the Glycoprotein_61-80_ peptide of LCMV at the C-terminus of the OmpC gene, alongside a Kanamycin resistance gene (CUMT8 GP61). *In vitro* culture of GP_61-80_- specific TCR transgenic T cells with heat-killed CUMT8 GP61 and splenic CD11c^+^ dendritic cells induced substantial proliferation not seen with stimulation with control CUMT8 strains (**Figure S4B**). To test whether neonatal *Enterobacteriaceae* colonization induces microbiota- specific Th17 responses we adoptively transferred congenic (CD45.1^+^ 10^5^ T cells/mouse) GP_61- 80_-specific TCR transgenic T cells into DOL10 pups (CD45.2^+^) and one day later colonized the mice with CUMT8 GP61 (**Figure 4C**). CUMT8 colonization of each pup was confirmed by culture of stool samples on Kanamycin plates (data not shown). Two weeks following the adoptive T cell transfer (DOL24), we analyzed siLP and mLN samples for activated GP_61-80_- specific CD4^+^CD45.1^+^ cells using flow cytometry (**Figure 4D**) and observed that in the absence of matIgA there was a substantial increase in the frequency and numbers of activated (CD44^hi^) GP_61-80_-specific TCR transgenic T cells in the mLN and sILP of pups fed by IgA deficient dams compared to controls, from which very few activated (CD44^hi^) cells could be recovered from any tissue (**Figure 4E-F**). In the absence of matIgA, GP_61-80_-specific TCR transgenic T cells predominantly differentiated into RORγt^+^FOXP3^-^ effector T cells and RORγt^+^FOXP3^+^ Tregs (**Figure 4G-H**). To observe whether microbiota-specific T cells could survive into adulthood, we assessed GP_61-80_ specific TCR transgenic T cell responses at DOL42 (6 weeks of age). The adult offspring of IgA^-/-^ dams had substantially increased populations of CD44^hi^ activated GP_61-80_ specific TCR transgenic T cells than C57BL/6 nursed mice (**Figure 4I-J**) that were similarly dominated by RORγt^+^ effector T cells and Tregs (**Figure 4K-L**). In concert with our microbiota and T cell analysis, as mice wean, the differences in colonization levels of CUMT8 GP61 (**Figure 4M**), and total Th17 cells (**Figure S4C-D**) equalize, indicating the increased survival of T cells activated in the adult offspring of IgA^-/-^ dams is not due to increased amount of persisting antigen. Taken together, we have demonstrated that a lack of matIgA during the pre-weaning period leads to the increased activation and long-term survival of *Enterobacteriaceae*-specific CD4^+^ RORγt^+^ T cells in the small intestine.

## Discussion

There are many long-term health benefits of breastfeeding over formula feeding and individuals who are breastfed are less likely to develop multiple autoinflammatory disorders(*55, 56*).

Modifications of the early life microbiome and immune system associated with modern lifestyles have also been implicated in the worldwide increase in autoinflammatory disease(*3*). Here we provide a link between breast milk antibodies and long-term predisposition to inflammation via the regulation of the neonatal microbiota and intestinal immune response.

Mammals evolved milk-feeding so that they could provide their infants complex nutrients at a time of their development when the infant cannot easily acquire these on their own. Milk also serves as a conduit for the mother to transmit immunological memory into the infant in the form of antibodies. These antibodies serve at least two main functions. First, they can be critical in protecting against enteric pathogens, which can cause lethal dehydration in infants. Secondly, as we have seen here, antibodies assist in the assembly and maturation of the small intestinal microbiota, hastening the replacement of facultative anaerobes, such as *Enterobacteriaceae*, with beneficial *Lactobacilli and Bifidobacteria.* It should be noted that a healthy complex microbiota also helps resist infection via colonization resistance and thus both of these mechanisms may have been primarily efforts to limit neonatal infection. The small intestinal microbiota is also more critical to the development of mucosal immunity because this tissue has a reduced and inconsistent mucus layer and numerous Peyer’s Patches that act as conduits for intestinal antigen transport and B cell activation(*57*).

The mechanism underlying why a lack of matIgA is leading to the development of long-lived *Enterobacteriaceae*-specific RORγt^+^ Th17 cells and Tregs is not entirely clear. One hypothesis consistent with the data is that matIgA is a critical component of a neonatal effort to ‘blind’ the immune response to early intestinal colonizers(*3*). The neonatal microbiota is both temporary and volatile, neonatal diet (milk) has very little antigenic diversity and the neonatal immune system lacks antigen-experienced memory cells. Thus, in the neonate there is not the same need to balance immunoregulation for the commensal microbiota with vigilance against pathogenic microorganisms and the default may be to respond to try to block and delay immunity to luminal antigens till after weaning. Indeed, the neonatal intestine is dominated directly after birth by *Enterobacteriaceae*, a family that contains many pathogens that could be lethal to infants and thus it is perhaps unsurprising that these bacteria are preferentially bound by maternal IgA and induce inflammatory effector T cell responses. Unfortunately, this predisposition to effector responses in the small intestine can create memory T cells that can be ‘recalled’ at later timepoints and contribute to inflammation. Future longitudinal study in human patients will be needed to determine if regulation of neonatal microbiota-responsive T cells is important to control the development of lifelong autoinflammation in genetically susceptible individuals.

## Acknowledgements

The authors would like to thank Y. Belkaid (NIH/NIAID) for providing IgA -/- mice and Mark Shlomchik (Univ. of Pittsburgh) for providing IgMi mice. We would also like to thank Susan Gottesman and Nadim Majdalani (NIH/NCI) for guidance in lambda red recombineering for the production of gp61-80 expressing CUMT8. The authors would like to thank the the Department of Pediatrics, and the Department of Immunology at the University of Pittsburgh for facilities and funding, as well as the members of the Hand, Poholek, Alcorn labs and A. Overacre- Delgoffe for helpful discussions. We would like to thank the staff of the Division of Laboratory Animal Services for animal husbandry, the University of Pittsburgh Health Sciences Sequencing Core at UPMC Children’s Hospital of Pittsburgh for RNA extraction and sequencing, the Center for Biological Imaging for training and use of the confocal microscope used in this study.

## Author Contributions

D.A.A., and T.W.H., designed the experiments; D.A.A., A.T.R., Y.C., D.C. and J.D.S. performed the experiments; D.A.A. and A.Y. analyzed the data; S.F., A.J.F. and T.W.H. created CUMT8 GP61; A.Y. and A.C.P. assisted in the design and analysis of RNA sequencing experiments; D.A.A. and T.W.H. wrote the manuscript.

## Materials and Methods

### Mice

Igha +/- mice were generated by pairing age matched C57BL/6 Jackson with Igha -/- originally provided by Y. Belkaid (NIH/NIAID). Comparisons between Igha +/- animals occurred between littermate matched breeding pairs. IgMi mice were generously provided by Mark Shlomchik (University of Pittsburgh)(*40*). Rag1-/-, and SMARTA-1 GP_61-80_ TCR transgenic T cells mice were acquired from Jackson Laboratories. All mice were maintained and all experiments were performed in an American Association for the Accreditation of Laboratory Animal Care-accredited animal facility at the University of Pittsburgh and housed in accordance with the procedures outlined in the Guide for the Care and Use of Laboratory Animals under an animal study proposal approved by the Institutional Animal Care and Use Committee of the University of Pittsburgh. Mice were housed in specific pathogen-free conditions unless otherwise noted.

### Microbial Strain

CUMT8 *E. coli* (CUMT8) was isolated by Kenneth Simpson (Cornell University, Ithaca NY)(*58*). CUMT8 *E. coli* was colonized with plasmid pSIM6 (S. Gottesman NIH/NCI) and electroporated with a geneBlock (IDT DNA) that contained the sequence of GP_61-80_ and a Kanamycin resistance gene targeted to the C-teminus/3’ UTR of the *E. coli* OmpC gene. For all transfers, CUMT8 was cultured overnight at 37°C in 5mL of Luria Broth from a frozen glycerol stock.

## Methods

### In vivo models

Gnotobiotic TIC Transfer – Total ileal contents (TIC) from DOL 14 IgA+/- pups fed by either a C57BL/6 or IgA-/- dam were isolated under aseptic conditions, pooled (≥3 littermate) and suspended in 2mL sterile PBS before passing through a 40μm filter. 200μL of TIC solution was gavaged into 6 week male gnotobiotic mice housed separately by group (IgA+/- TIC fed by either a C57BL/6 or IgA-/- dam) and maintained in germ free conditions on standard chow. Mice were taken down 10 days following gavage for small intestine lamina propria (siLP), Peyer’s patch (PP), and mesenteric lymph node (mLN) flow cytometric analysis.

### Antibiotic Treatment

Ciprofloxacin hydrochloride monohydrate (Fisher Scientific) was dissolved in sterile water and filtered through a 0.2μm filter, for a final solution of 6mg/mL. IgA+/- pups were gavaged daily with 30μL of solution on DOL12-15, for a daily dose of 0.18mg/day(*59*). On DOL24 pups were harvested and siLP analyzed.

### Injury model

To induce intestinal damage in adult mice, we gavaged indomethacin (SigmaAldrich) daily for three days (6.25mg/kg/day). We weighed the animals at D0-D3, and harvested small intestine and mLN onD3 for either histology or tissue preparation for flow cytometry. For CD4 depletion during indomethacin treatment, we gave intraparitoneal injections of anti-CD4 antibody (BioXCell. clone: GK1.5) (200μg) every three days, starting at Day(-3).

### SMARTA GP_61-80_ TCR transgenic T cell adoptive transfer

Spleen and mLN were harvested from naïve CD45.1+ SMARTA mice (Jackson Labs) and processed into a single cell suspension. After lysis of red blood cells using ACK lysis buffer (Thermo Fisher) following manufacturers instructions, we enriched for CD4 cells using negative selection with CD19 and CD11c. To do this, we stained spleen + mLN with CD19 PECF594 and CD11c PECF594 and performed magnetic selection with EasySep PE positive selection kit (Stem Cell Technologies) following manufacturer’s instructions. We then stained the negative fraction, and sorted on a BD FACS Aria for live CD4+CD44-CD25-CD45.1+Vα2+. Sorted fraction was checked for purity and counted using a Cellometer (Nexcelom) with AOPI (Viastain Nexcelom) staining to validate cell quantity, and resuspended in sterile PBS at a concentration of 2x10^6^ cells/mL to transfer 1x10^5^ cells/50μL per DOL9-11 pup via intraperitoneal injection.

### Preparation CUMT8

Following overnight growth in Luria Broth at 37°C, we quantified CUMT8 growth by OD_600_ reading on a spectrophotometer, and calculated CFU using an online OD_600_ converter (Agilent). We then spun down the appropriate volume of culture and resuspended in sterile PBS to a concentration of 1x10^8^ colony forming units (CFU)/mL for a final inoculation of 8x10^6^ CFU/50μL per pup.

### Bacterial Flow Cytometry

Stool or TIC samples were weighed and suspended in PBS. Following a low speed spin (50xg 5 minutes) to remove debris, bacterial supernatant was washed 1x with PBS and spun at 5000xg for 5 minutes. For quantification of antibody binding, bacteria were stained with listed concentrations of antibodies in key resource table. All antibody staining was carried out in PBS containing 10% normal rat serum for 30 minutes on ice, followed by 3x PBS washes (7000xg 5 minutes). Bacterial suspension was fixed with 4% paraformaldehyde in PBS for 15 minutes at 22C, followed by 3x PBS washes. For fluorescence in situ hybridization (FISH), fixed samples were resuspended in 200ul FISH hybridization buffer prewarmed to 50C containing appropriate probes at a concentration of 100nmol. Following overnight incubation in a shaking heat block at 50C, samples were washed 3 times (10 minute incubation in 1mL prewarmed FISH wash buffer, spun down at 7000xg 5 minutes). All bacteria, antibody stained or FISH, were counterstained with Hoescht 33342, in PBS or FISH wash buffer for 5 minutes, followed by 3 washes (1mL PBS/FISH wash buffer 7000xg 5min). Immediately prior to running on BD Fortessa flow cytometer, samples were resuspended in 500ul PBS and filtered through 40μm filter paper. For quantification, we utilized CountBright absolute counting beads (Invitrogen) following manufacturer’s instructions.

### Preparation of tissue sections for imaging

Sections of small intestine with luminal contents were fixed with methacarn (60% methanol, 30% chloroform, 10% glacial acetic acid) overnight at 4C and washed with 70% EtOH before being sent for paraffin embedding and sectioning. 5μm paraffin sections were deparaffined with successive washes of Xylene (2x 8 min), 95% EtoH (1x 10min), 90% EtOH (1x 10min), diH2O (1x 10min).

### Immunofluorescence

Deparaffined sections were blocked with 10% normal rat serum 0.5% Tween-20 in PBS for 1hr at 4C. Following 3x washes with PBS, samples were stained with conjugated antibodies in 1%

normal rat serum + 0.5% Tween-20 in PBS overnight at 4C. Tissues were washed 3x with PBS, and stained with Hoescht for 5 minutes at room temperature, followed by mounting with prolong antifade mountant (ThermoFisher). Slides cured overnight at 4C before staining on a Nikon NI-E confocal microscope at the Pittsburgh Center for Biological Imaging.

### Preparation of single cell suspensions from mouse tissues for flow cytometry

Small intestine and mesenteric lymph nodes (mLN) were harvested in 3% complete RPMI. mLN was pressed through a 70μm filter with a syringe and washed with complete media to acquire single cell suspension. Small intestine was processed to remove fat and connective tissue, and Peyer’s patches were removed by surgical scissor and placed in 3% complete media. Acquisition of lymphocytes from siLP and intraepithelial lymphocytes (IEL) was performed as previously described. Peyer’s patches lymphocytes were isolated by incubation of tissue in 10mL 0% complete media containing 1mg/mL Liberase DL and 1mg/mL DNAse at 37°C for 25minutes.

Digestion reaction was quenched with the addition of an equal volume of 3% complete media, and suspensions were filtered through first a 70μm, then a 40μm filter. For transcription factor analysis, single cells suspensions were stained with fixable live dead aqua (Thermo Fisher), fixed and stained with transcription factor staining buffer set (eBioscience) following manufacturers instructions. For cytokine expression, single cell suspensions were plated in complete media with 10% FBS for 4hrs with brefeldin A, Phorbol-12-myristate-13-acetate (PMA), and Ionomycin followed by staining with fixable live dead aqua (Thermo Fisher), fixed and stained with cytofix/cytoperm (BD Biosciences) following manufacturer’s instructions.

## QUANTIFICATION AND STATISTICAL ANALYSIS

### 16S#rRNA Microbiome analysis

Total ileal contents were extracted from IgA+/- pups from littermate matched breeding pairs. For each timepoint, samples were taken from at least 2 separate cages to control for cage variation of the microbiome and stored at -80C until acquisition of all sample groups. We extracted DNA from total ileal contents using QIAmp Fast DNA stool kit (Qiagen) following manufacturer’s instructions, and quantified DNA content by Qubit 4 fluorometer (Thermo Fisher). We sent DNA for 16S ribosomal sequencing (Microbiome Insights Inc.). Analysis of microbiome sequences was carried out using Qiime2(*60*). Raw sequences were filtered and denoised using DADA281. We aligned amplicon variant sequences (ASVs) using mafft, and constructed a phylogeny with fasttree2. Samples were rarefied to 63,000 sequences per sample before Alpha diversity metrics (Faith’s phylogenetic diversity and Pielou’s evenness) were assessed. We aligned ASV’s taxonomy using naïve Bayes taxonomy classifier (greengenes 18_9 99% OTU’s) and utilized ANCOM to compare order level abundances between groups.

### Epithelial RNAsequencing

Small intestine was removed from DOL 14, DOL 24, and DOL 42 IgA+/- mice fed by either a B6 or IgA-/- dam. For DOL 42 timepoint, mice were cohoused with animals of the same gender following weaning. Small intestine epithelial cells were isolated using a protocol modified from a previous study(*61*). Small intestine was butterflied and rinsed with PBS to remove luminal contents. Using cut tissue into 1cm sections and placed in 20mM EDTA in PBS on ice for 60 minutes, shaking vigorously every 20 minutes. We performed a slow speed spin (50xg 5min) to pellet large tissue sections, and removed supernatant to a new 50mL tube. We repeated the same process two more times, with 30 minute incubations of tissue sections on ice. To confirm successful extraction of small intestinal crypts from the tissue, we looked at each supernatant fraction under 20x magnification on a lightfield microscope. Following the final incubation step, tissue sections were discarded and the combined supernatant fractions were spun down at 1650xg for 10 minutes at 4C, and resuspended in 10mL of PBS to wash. Wash step was repeated 2x times. We resuspended epithelial pellet in TrypLE express (Invitrogen) at 37°C for 5min. We quenched the TrypLE express with a 5x volume of RPMI+ FBS and filtered through a 40μm filter. We quantified epithelial cells using a Cellometer (Nexcelom) and AO/PI live/dead (Nexcelom) discrimination.

To acquire purified epithelial cells, we stained and sorted (FACS Aria - BD Biosciences) single cell suspension for Live EpCAM+CD45-Ter119- and quantified sorted cells with a cellometer (Nexcelom) and AO/PI live/dead (Nexcelom) discrimination. We spun down cells and flash froze the pellet on dry ice before storing at -80C.

### RNA-seq Library Generation and Sequencing

RNA was assessed for quality using an Agilent TapeStation 4150 and RNA concentration was quantified on a Qubit FLEX fluorometer. Libraries were generated with the Takara SMART-Seq Stranded kit (Takara: 634447) using Takara SMARTer RNA Unique Dual Index A and B Kits (Takara: 634452, 634457) according to the manufacturer’s instructions. RNA was normalized to 10 ng/µl in a total volume of 7μl of input RNA. RNA with RIN scores ≥4 were fragmented for 4 minutes, RNA with RIN scores <4 were not fragmented. 5 cycles were used for PCR1, followed by ribosomal RNA depletion using scZapR. No samples were pooled prior to PCR2 where 12 cycles were completed. Library assessment and quantification was done using Qubit 1x HS DNA assay kit on a Qubit 4 fluorometer, and a HS NGS Fragment kit on an Agilent 5300 Fragment Analyzer. Libraries were normalized and pooled to 4nM by calculating the concentration based off the fragment size (base pairs) and the concentration (ng/µl) of the libraries. Sequencing was performed on an Illumina NextSeq 2000, using a P2 100 flow cell for an average of 25 million reads per sample. The pooled library was loaded at 850 pM. Sequencing was carried out 2x59 bp, with a custom run chemistry utilizing 3 dark cycles at the start of R2 that encompasses the 3 bp of the Takara adapter present in the read.

Library generation and sequencing was performed by the University of Pittsburgh Health Sciences Sequencing Core (HSSC), Rangos Research Center, UPMC Children’s Hospital of Pittsburgh, Pittsburgh, Pennsylvania, United States of America.

### Sequencing Data Pre-Processing

FastQC was used to perform a quality assessment on all fastq files. The mouse reference genome (GRCm38) was downloaded from Ensembl. Adaptors were trimmed using cutadapt v1.18. RNA sequencing samples were aligned using HISAT2 v2.1.0. Raw count values were generated using Subread, and gene expression values were normalized using transcripts per million.

### Gene Expression Analysis

DESeq2 was applied to raw counts transcriptome data in a pairwise manner to determine differentially expressed genes. These gene lists were compiled to make a master list of all differentially expressed genes. Log2 transformed normalized counts were then displayed as a heatmap to display expression of differentially expressed genes.

Gene set enrichment analysis Small intestinal epithelial cell gene signatures were acquired from a single cell dataset and input for comparison to experimental epithelial gene set using GSEA. GSEA analysis was carried out with GSEA v4.3.1 obtained from www.broadinstitute.org. Enrichment of selected gene set was characterized by significance of enrichment score and false discovery rate (FDR).

### CUMT8 quantification

Stool pellets from animals gavaged with CUMT8 were collected weekly and weighed prior to DNA extraction using QIAmp Fast DNA Stool mini kit following manufacturers instructions. CUMT8 was quantified using primers for a highly conserved 16S region, and a previously published primer set for CUMT8 FliC3(*62*). To determine relative quantity of CUMT8 we took the inverse of average Ct value of three technical replicates and divided by the weight of the original stool pellet.

### Imaging quantification

Enteroendocrine cells were defined as chromogranin A + cells within the small intestine epithelium. Images were analyzed using ImageJ, and brightness contrast adjusted to remove background signal using an unstained control slide. Chromogranin A+ cells were quantified per 5 continues villi with crypts visible on each slide, on 3 individual sections per each mouse, with the average of the three sections plotted.

### Statistics

Data are presented as mean + SEM unless otherwise noted. Listed n is the number of mice in the experiment, unless noted to be a compilation of multiple experiments. Unpaired students t test was used when comparing two groups, one way ANOVA used for multiple comparisons. For comparing weight loss curves we used simple linear regression analysis. Figure legends provide individual information on sample numbers and statistical analysis for individual experiments.

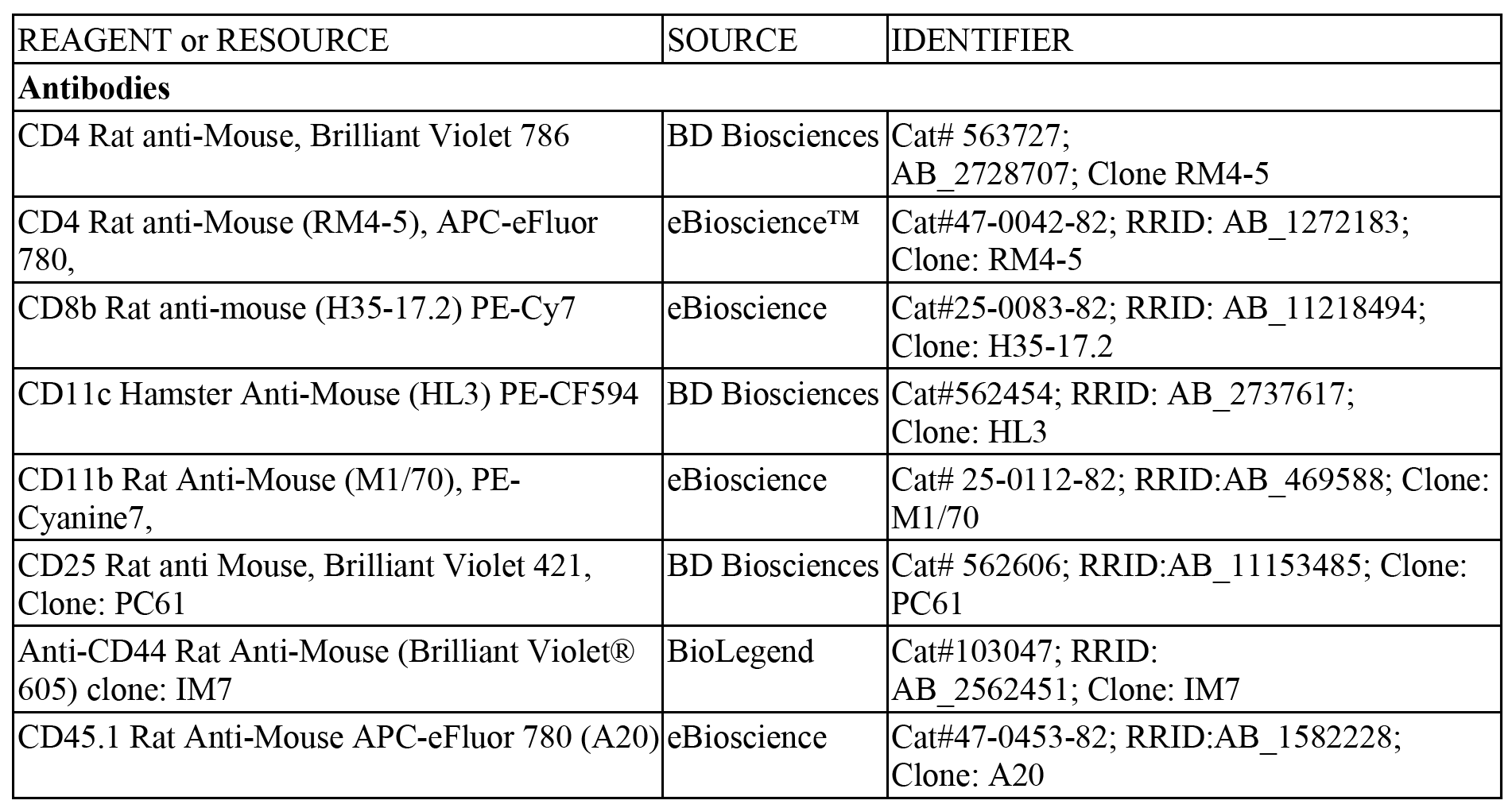
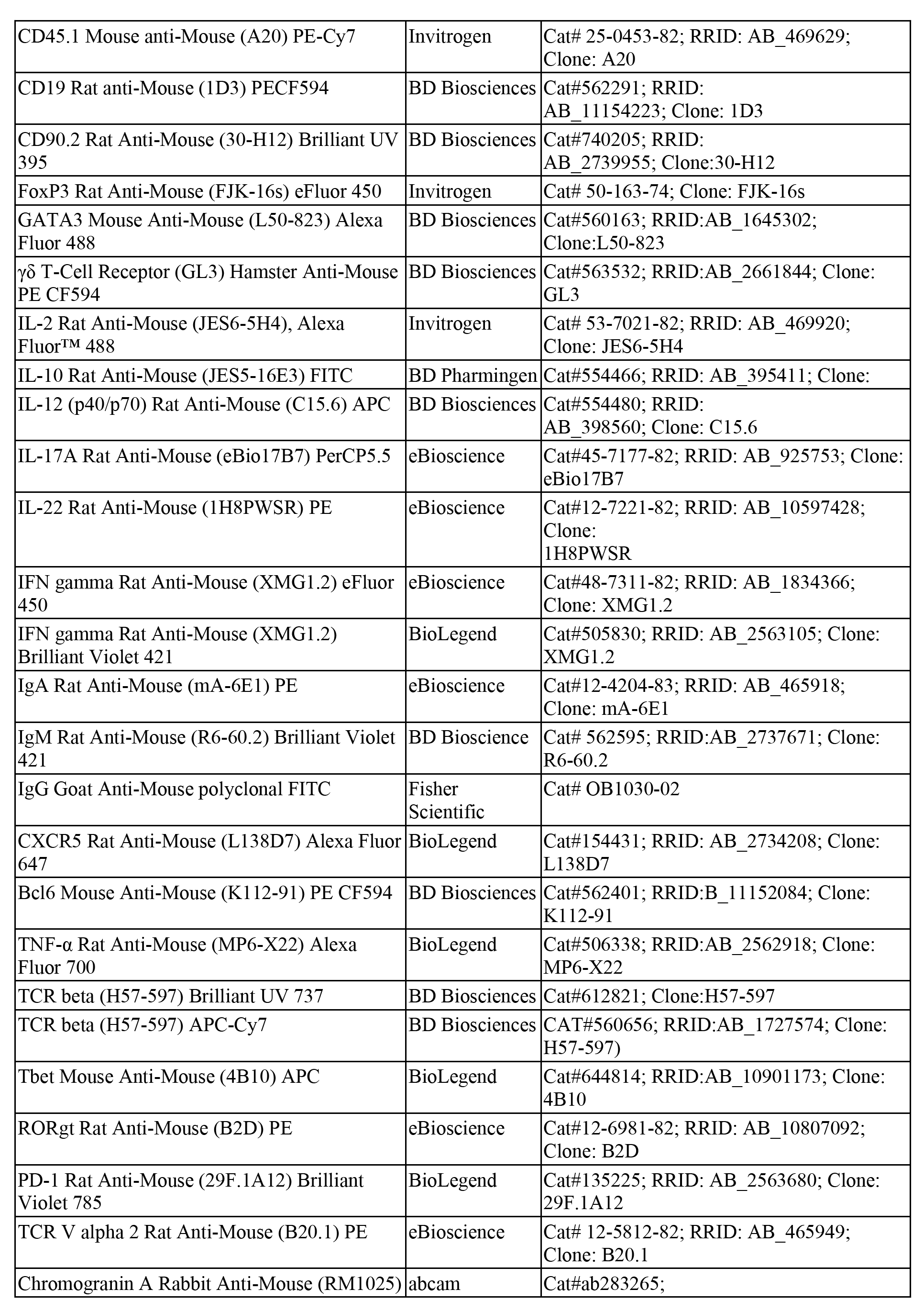
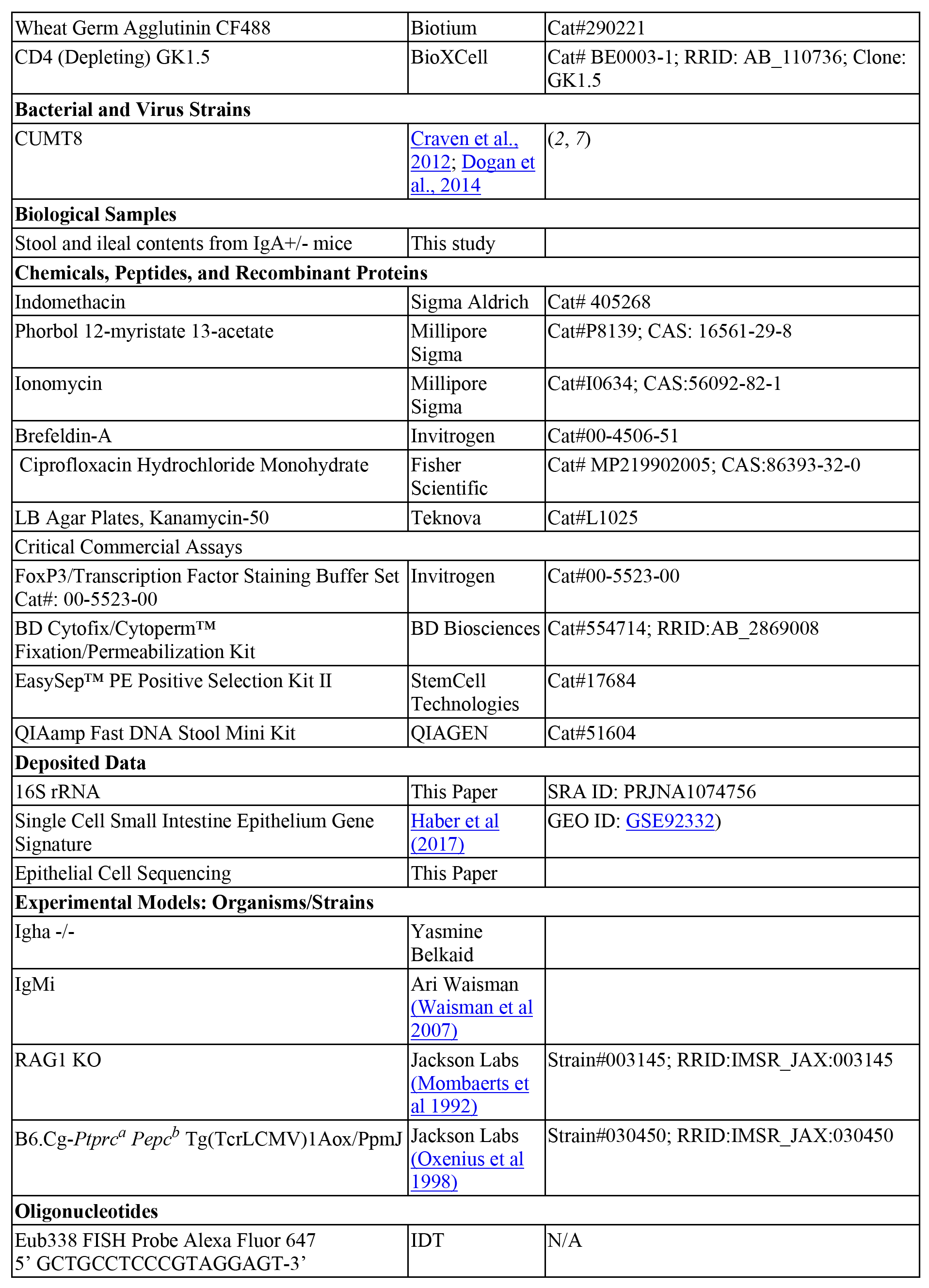
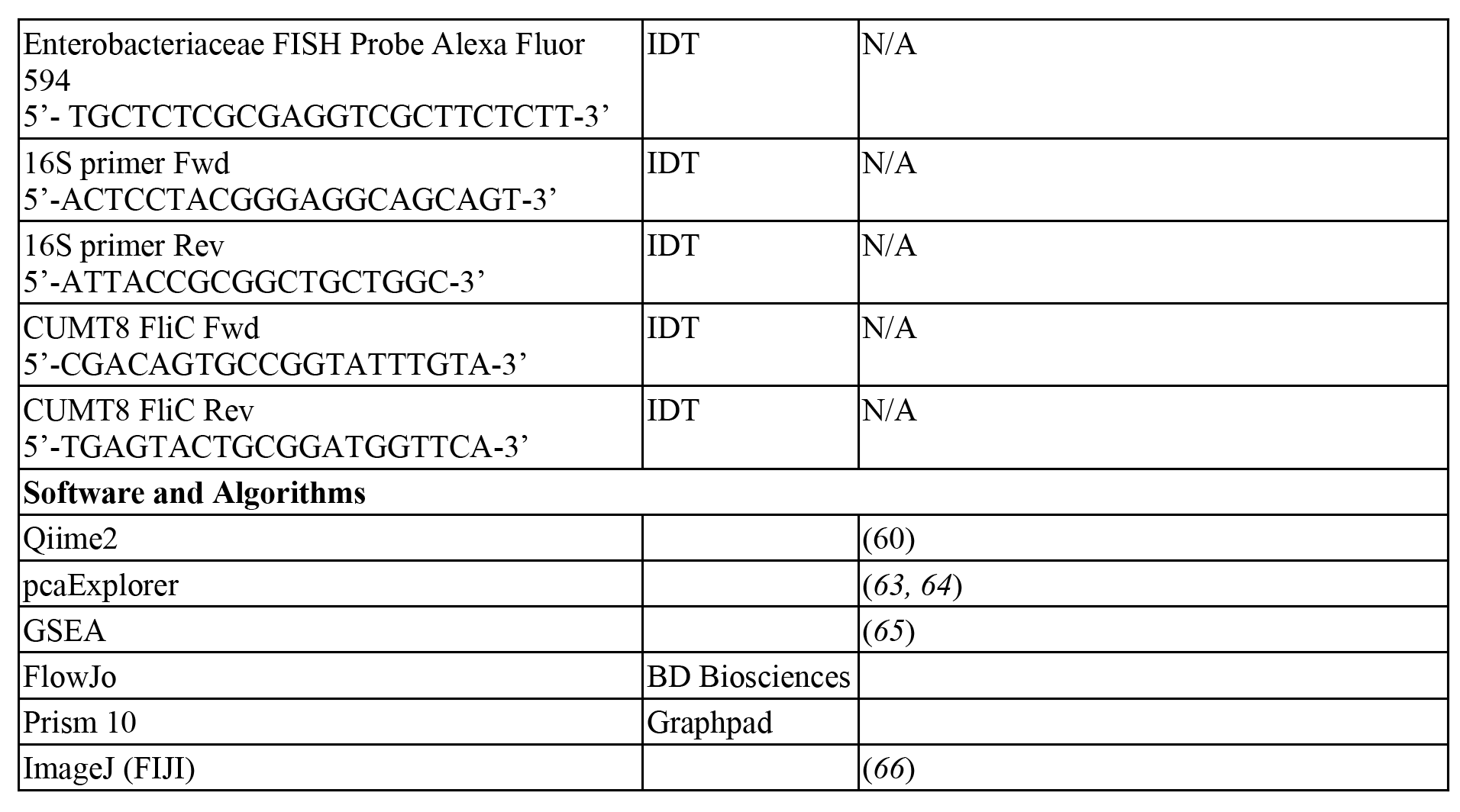

---

**Figure S1.**
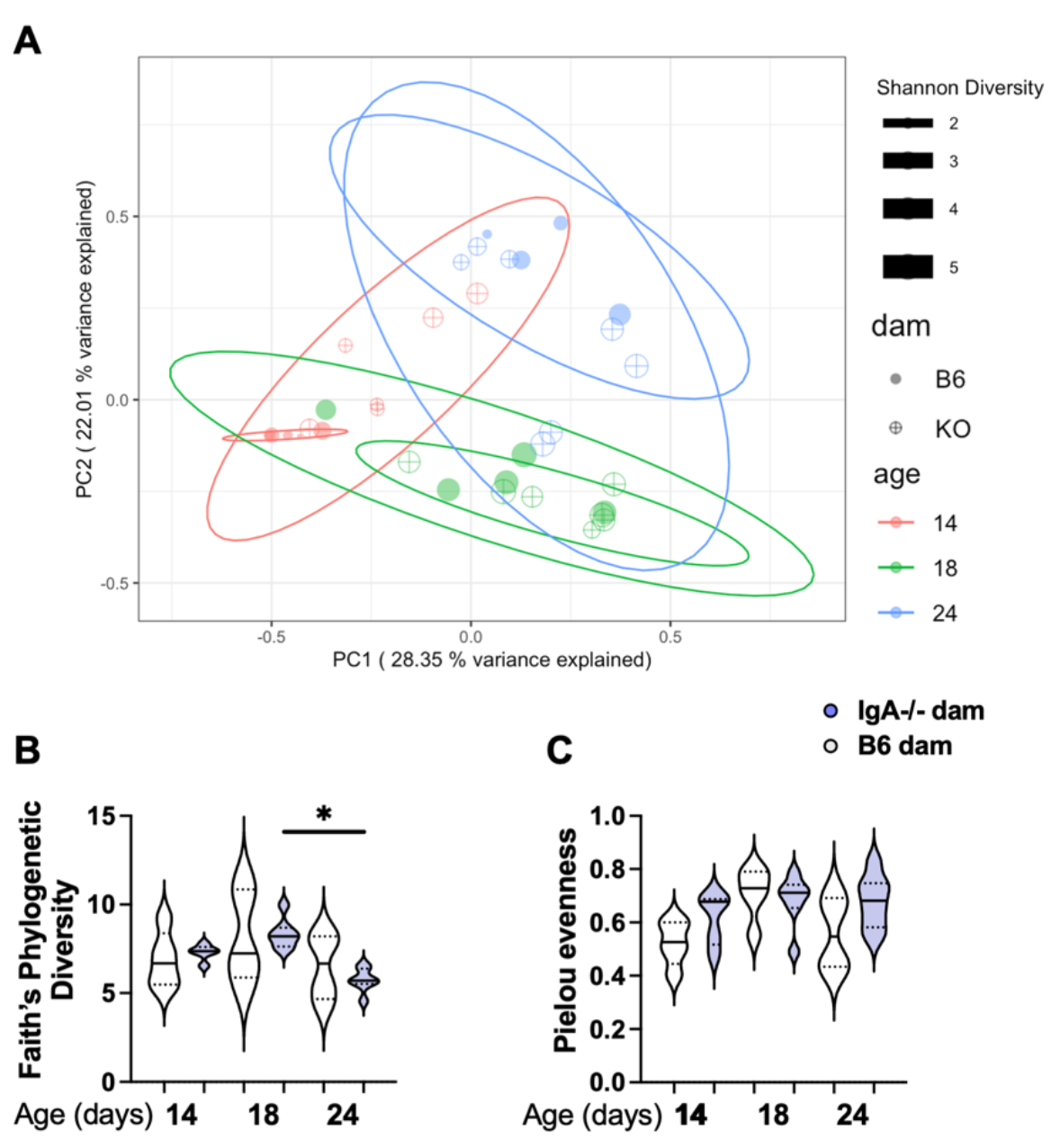
Maternal IgA induces transient changes to the small intestinal microbiome. Pups were bred as in Figure 1A (A) Genomic DNA isolated from TIC of pups analyzed by 16S rRNA gene amplicon sequencing. Shown is PCoA of DOL 14, 18, 24 in the presence or absence of matIgA. (Bray-Curtis). Each point is one mouse, and the size of the point represents Shannon diversity. Ellipse drawn for each group by 95% confidence level for a multivariate t-distribution and colored by pup age (DOL). (B-C) Alpha diversity of 16S rRNA sequences. (B) Faith’s phylogenetic diversity and C) Pielou’s evenness analysis. Significance for B-C assessed by one-way Anova with multiple comparisons. *p<0.05, **p<0.01, ***p<0.001.

**Figure S2.**
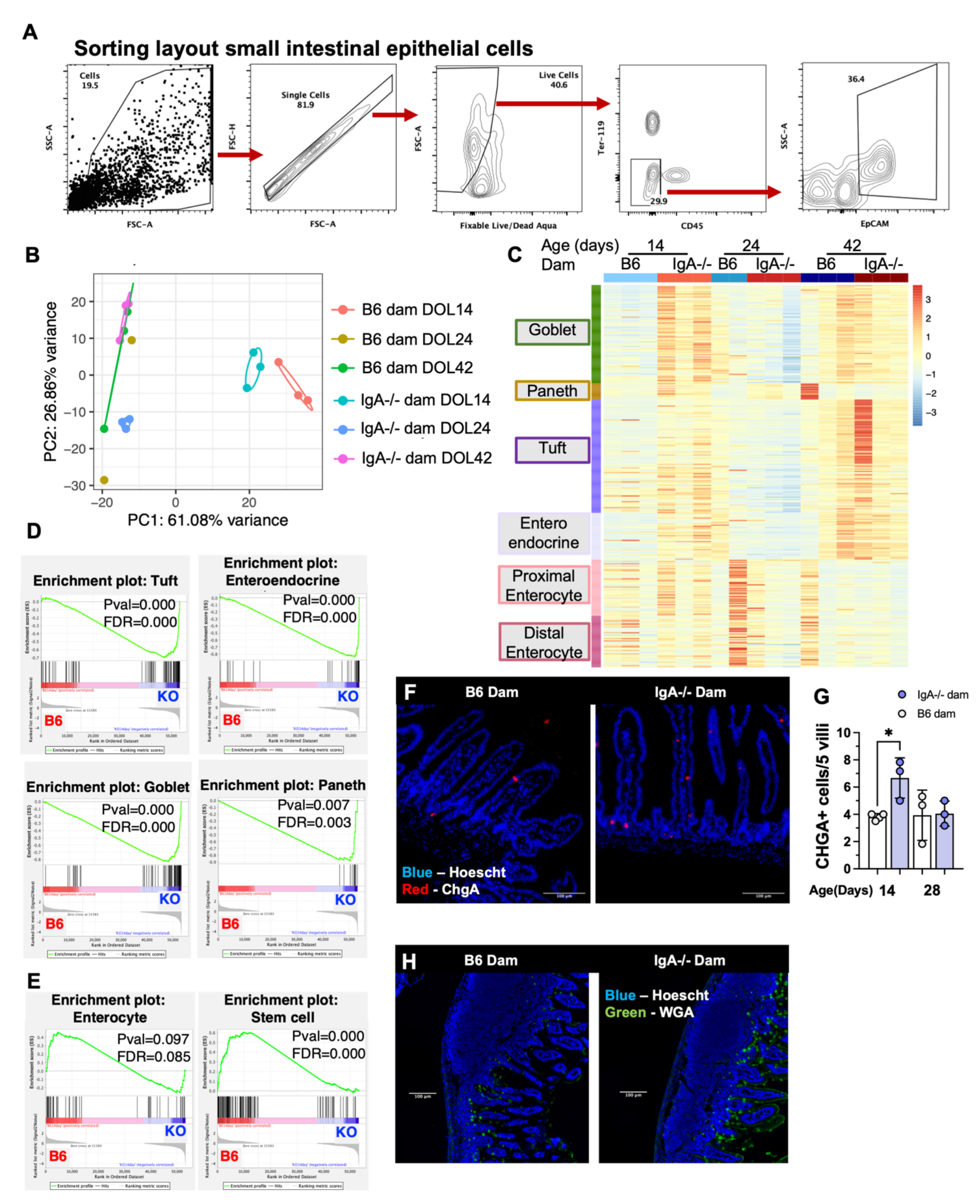
Development of the neonatal small intestinal epithelium is regulated by matIgA . . Pups were bred as in Figure 1A (A) Example of flow cytometric sort layout utilized to acquire small intestinal epithelial cells for RNA sequencing. (B-E) EpCAM+ cells were isolated from the siLP of pups nursed by either a IgA+/+ or IgA-/- dam at DOL 14, 24, and 42. RNA from all groups was extracted and sequenced. (B) PcoA plot with 0.5 confidence ellipses around indicated groups (created with pcaExplorer). (C) Heatmap of all groups expression of gene signatures associated with specific intestinal epithelial cell types. (D-E) GSEA enrichment analysis of epithelial cell type gene sets comparing pups nursed by IgA-/- and IgA+/+ dams. (D) Gene sets enriched in the pups of IgA-/- dams. (E) Gene sets enriched in the pups of IgA+/+ dams. (F) Paraffin embedded sections of the distal small intestine of DOL 14 pups fed by indicated dam were stained with anti-Chromogranin A (ChgA-red) and counterstained for nucleic acids (Hoescht -Blue). (G) Quantification of Chga+ cells/5 continuous villi. Quantification is the average of ChgA+ cells/5 villi. (H) Representative images of small intestines of pups nursed by the indicated dam at DOL14. Sections were stained with wheat germ agglutinin (WGA) to stain for mucus and mucus producing goblet cells and counterstained with Hoescht to detect nucleic acid. Images acquired at 20x. (D+E) Statistical analysis of gene enrichment calculated using GSEA analysis, displayed false-discovery rate (FDR) and nominal p-value. Unpaired student’s T test was used for (G). *p<0.05, **p<0.01, ***p<0.001. Each point is one mouse.

**Figure S3.**
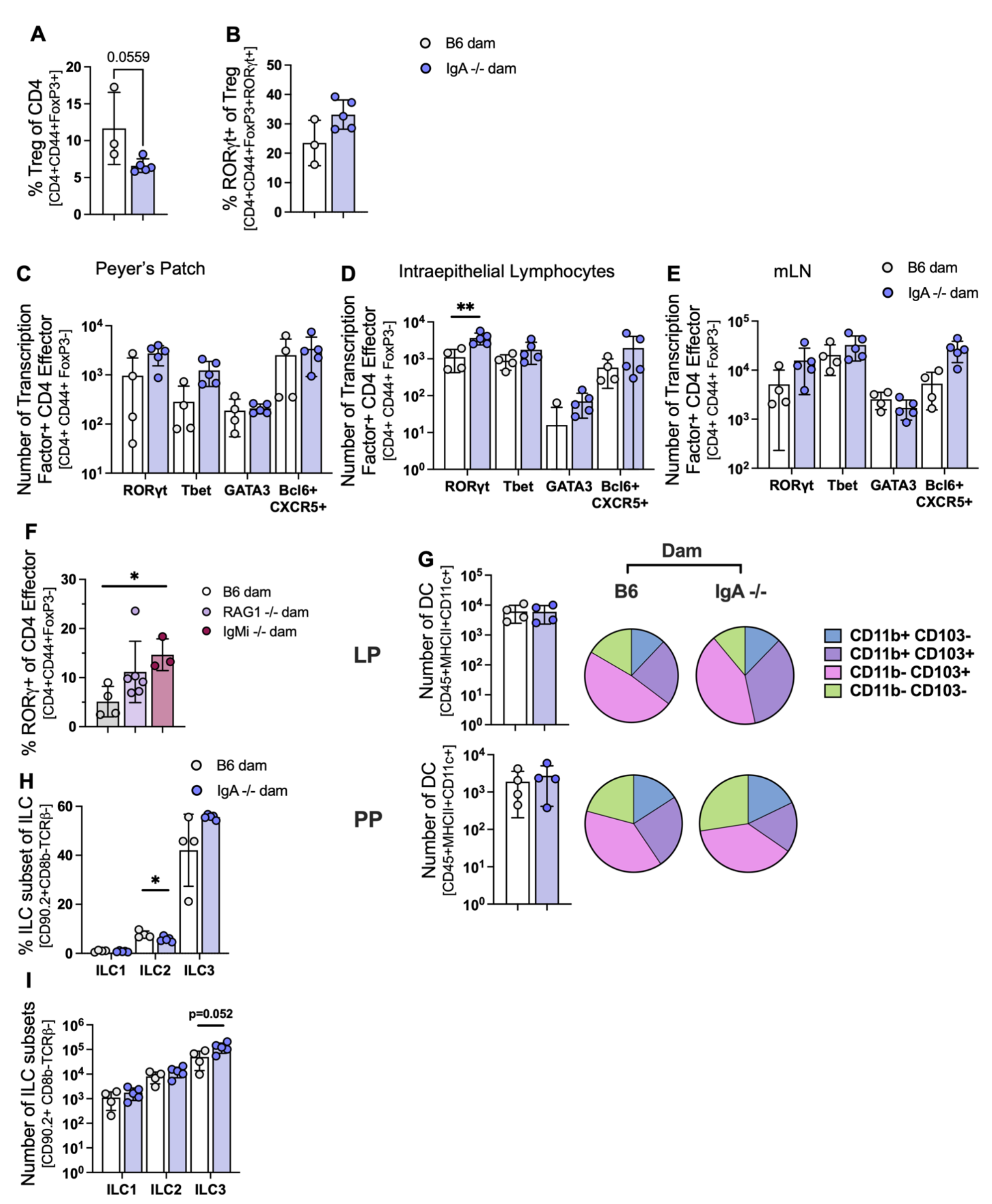
Immune cell characterization of the pups of IgA-/- or IgA+/+ dams. Pups were bred as in Figure 1A. (A-B) Analysis of siLP of DOL 24 IgA+/- pups. (A) Relative proportion of Tregs (FoxP3+) of CD4+ T cells (live, TCRβ+, CD90+, CD4+). B) Relative proportion of RORγt + Tregs (live, TCRβ+, CD90+, CD4+, FoxP3+). (C-E) Transcription factor expression profiling of CD4+ effector T cells (live, TCRβ+, CD90+, CD4+, FoxP3-, CD44+) within (C)Peyer’s patches, amongst (D) intraepithelial lymphocytes and in the (E) mesenteric lymph nodes. (F) Relative proportion of CD4+ effector T cells (Live TCRβ+CD4+CD90+Foxp3-) expressing RORγt+ in siLP of DOL 24 pups (IgA+) fed by either a Rag1-/- or IgMi dam. (G) Flow cytometric analysis of antigen presenting cells of DOL 14 IgA+/- pups. Measurement of dendritic cells (live, CD45+, MHCII+, CD64-, CD11c+ within Peyer’s patch (PP) and mesenteric lymph node (mLN) as labeled. Bar plot displays number of dendritic cells (DC) in each tissue, with the pie chart displaying the relative proportion of CD103 and CD11b expressing cells within the DC population. (H-I) Innate lymphoid cell (ILC) subsets in the siLP of DOL24 IgA+/- pups by (H) relative proportion of total ILC (Live, TCRβ-, CD8b-, CD90+) and (I) Total count. Subsets were characterized by transcription factor expression as follows; ILC1 (Tbet+), ILC2 (GATA3), ILC3(RORγt). Results are representative of 2-3 separate experiments. Unpaired student’s T test was used for all analysis. *p<0.05, **p<0.01, ***p<0.001. Each point is one mouse.

**Figure S4.**
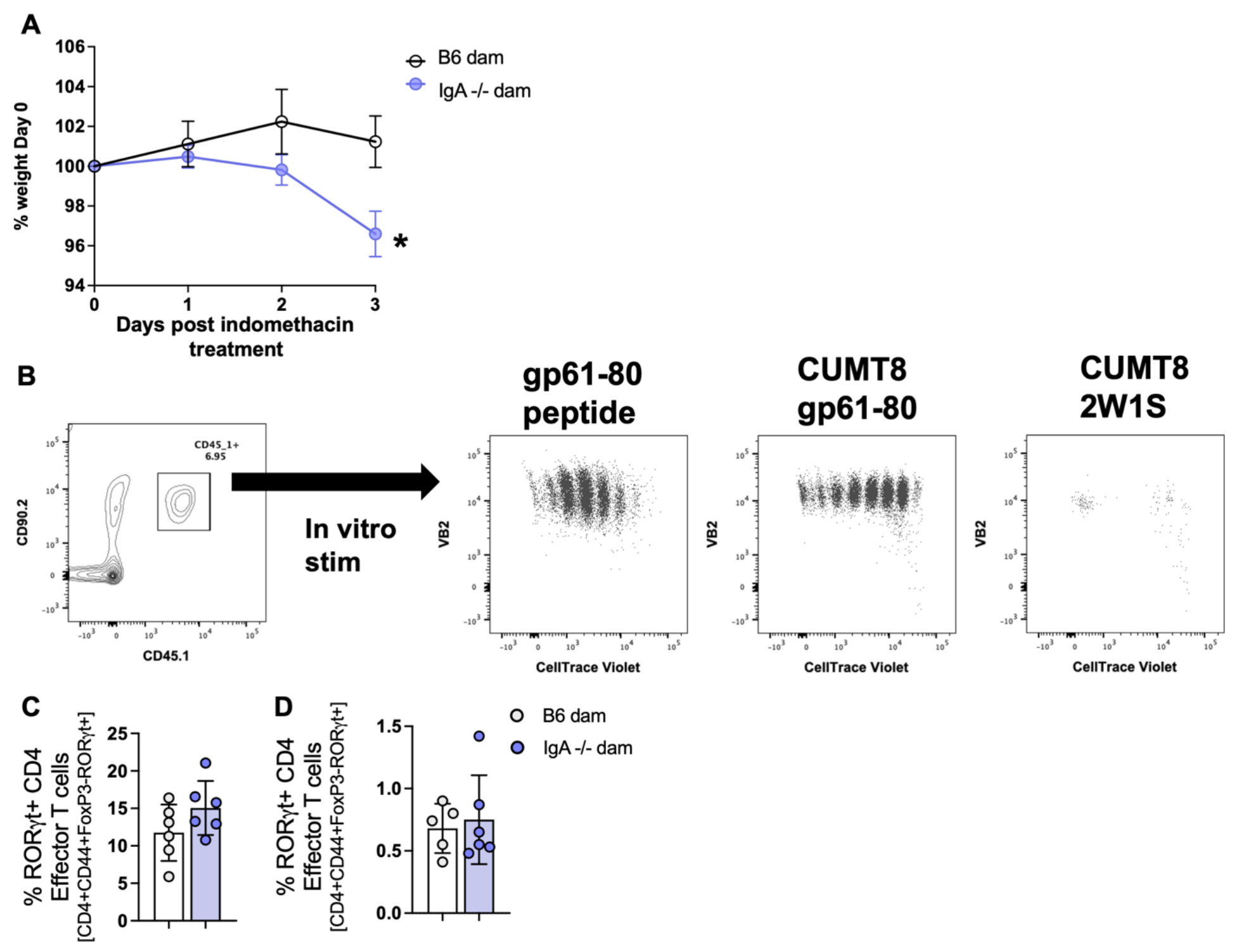
Long-term effects of maternal IgA deficiency on intestinal T cell populations. Pups were bred as in Figure 1A .(A) Pups nursed by either IgA-/- or IgA+/+ dams were cohoused following weaning until six weeks of age, at which point all groups were gavaged with 6.25mg/kg/day of indomethacin for three days. Weight change was monitored daily and calculated as % of weight at day 0, pooled from 2 independent experiments. (B) *In vitro* stimulation of GP_61-80_ TCR transgenic T cells prestained with Celltrace violet. Magnetically selected CD11c+ cells from a naïve C57BL/6 mouse were plated with indicated stimuli (GP_61-80_ peptide, heat killed CUMT8 expressing GP_61-80_ peptide, and heat killed CUMT8 expressing the irrelevant peptide 2W1S) for two hours at 37°C. Sorted GP_61-80_ TCR transgenic T cells (CD4+ CD44 low CD25-TCRVα2+) labeled with cell trace violet were added to the pre-loaded DC and incubated for four days at 37°C before flow cytometric analysis. (C-D) 1x10^5^ CD45.1+ naïve (CD44^lo^CD25-) GP_61-80_ TCR transgenic T cells were transferred by IP injection into pups at DOL10-11. CUMT8 GP_61-80_ was gavaged one day after transfer. (C) Flow cytometric analysis of the relative proportion of RORγt+ of CD4+ CD45.1- effector cells in the siLP (C) and mLN (D) at DOL42. (A) Simple linear regression was performed *p<0.05, **p<0.01, ***p<0.001. Students t test was used for C+D. Each point is one mouse.

**Table. S1.**
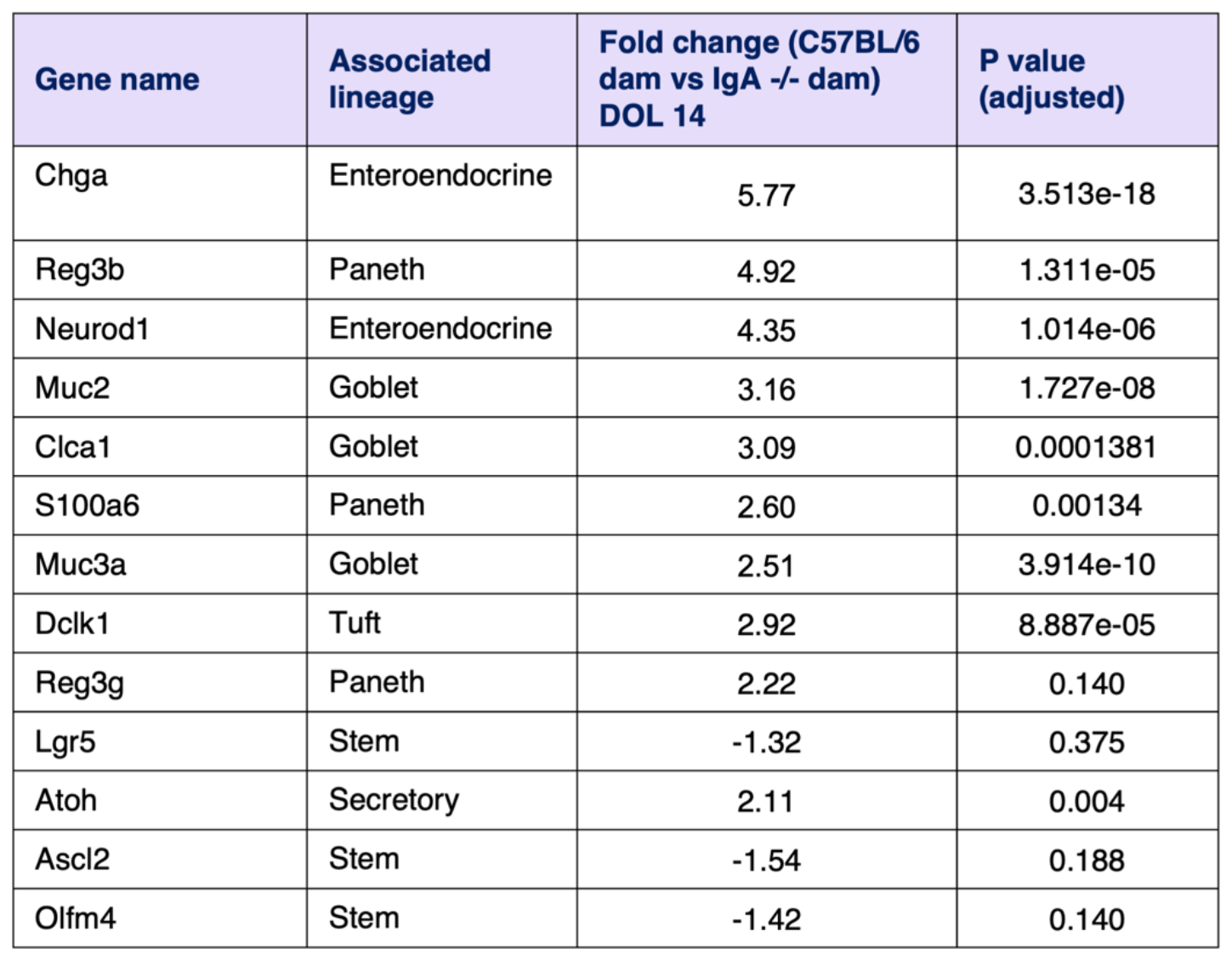
Selected genes from RNA sequencing analysis of small intestinal epithelial cells from day 14 IgA+/- pups.

| **Gene name** | **Associated lineage** | **Fold change (C57BL/6 dam vs IgA <sup>-/-</sup> dam) DOL 14** | **P value (adjusted)** |
| --- | --- | --- | --- |
| Chga | Enteroendocrine | 5.77 | 3.513e-18 |
| Reg3b | Paneth | 4.92 | 1.311e-05 |
| Neurod1 | Enteroendocrine | 4.35 | 1.014e-06 |
| Muc2 | Goblet | 3.16 | 1.727e-08 |
| Clca1 | Goblet | 3.09 | 0.0001381 |
| S100a6 | Paneth | 2.60 | 0.00134 |
| Muc3a | Goblet | 2.51 | 3.914e-10 |
| Dclk1 | Tuft | 2.92 | 8.887e-05 |
| Reg3g | Paneth | 2.22 | 0.140 |
| Lgr5 | Stem | -1.32 | 0.375 |
| Atoh | Secretory | 2.11 | 0.004 |
| Ascl2 | Stem | -1.54 | 0.188 |
| Olfm4 | Stem | -1.42 | 0.140 |

